# Examining the fitness benefits of social plasticity to prey availability in bottlenose dolphins

**DOI:** 10.1101/2025.08.21.671441

**Authors:** David N. Fisher, Barbara J. Cheney

## Abstract

A key explanation for phenotypic plasticity is that changing behaviour to match conditions increases fitness. Affiliative social associations with conspecifics can be important for coping with challenging conditions, and therefore plasticity in social behaviour in response to environment variation may bring fitness benefits. Here we leverage a dataset of 711 observations of 62 female bottlenose dolphins (*Tursiops truncatus*) over 35 years to estimate the plastic responses in social behaviour to salmon (*Salmo salar*) abundance and determine if either this plasticity or their mean social behaviour is associated with lifetime calving success. We found no evidence that mean social behaviour or plasticity in response to salmon abundance were associated with this component of fitness. Female dolphins did not seem to adjust their social behaviour to buffer low, or take advantage of high, salmon abundance and this prey variability was not associated with female reproductive success. Limitations in the data and follow-up simulations suggest that these null relationships, especially for plasticity, may be due to a lack of power, highlighting the difficulty of estimating these covariances even in long term studies with large datasets.

## Introduction

Animals are frequently exposed to adverse environmental conditions, including extreme weather events or fluctuations in prey availability and predation risk, which can impact their fitness. To mitigate these factors, animals may change their traits (show phenotypic plasticity) to buffer any fitness costs that may otherwise be inevitable in adverse conditions (Blumstein et al., 2023; Candolin & Wong, 2011; López-Sepulcre & Kokko, 2012). For instance, brown trout *Salmo trutta* may decrease their metabolism at times of low food availability (Auer et al., 2015), while goldcrest *Regulus regulus* form larger groups to huddle and conserve heat in cold conditions (Hogstad, 1984). Behaviours are the type of trait that are often thought to be animals’ first line of defence against adverse conditions, as they can change relatively quickly compared to morphological or even physiological traits (Sih, 2013; Sih et al., 2011). Consequently, changing behaviour to ameliorate the costs of environmental stressors is thought to be widespread (Tuomainen & Candolin, 2010; Wong & Candolin, 2015).

Many animals live in groups or near others, and so engage in social behaviours; interactions with conspecifics that can be cooperative or competitive in nature (Rubenstein & Abbot, 2017; Székely et al., 2010). Social behaviours are therefore a key mechanism through which animals may be able to buffer against environmental change (Komdeur & Ma, 2021). Global-scale comparative analyses have been used to test the general idea that social interactions with conspecifics helps animals cope with harsh environments. These relate aspects of a species’ social structure, such as the presence or absence of cooperative breeding, to environmental parameters of the species’ native range (e.g. variability in rainfall). For example, higher incidences of communal living are found in social spiders in areas with higher rain intensity which challenge the survival of lone females (Guevara & Avilés, 2015). These analyses have demonstrated a link between communal and cooperative breeding systems and survival in harsh conditions, supporting the idea that social living helps animals cope with environmental stressors (Firman et al., 2020; Guevara & Avilés, 2015; Jetz & Rubenstein, 2011; Sheehan et al., 2015).

Comparative analyses, however, do not shed light on whether *plasticity* in behaviour helps animals survive and reproduce in adverse conditions. Such studies require multiple measures of individual social behaviour across a range of environments and to associate variation in the social behaviours with a fitness component. Recent studies have demonstrated a clear interaction between social behaviour or social conditions (such as group size) and environmental factors in a regression on fitness components like survival or reproductive success (Borger et al., 2023; Guindre-Parker & Rubenstein, 2020). This suggests that animals can adjust their social behaviours to limit the harm caused by stressful environments, and the authors argue this provides evidence for adaptive social plasticity.

However, while the analyses conducted by Borger, Guindre-Parker, and colleagues shows whether particular behaviours of individuals bring benefits in certain conditions and not others, they do not explicitly test whether plasticity itself is beneficial. Distinct subsets of individuals could show different behaviours when environmental conditions are compared, which would still result in a clear statistical interaction. Instead, as proposed by Warrington et al. (2024), a more appropriate test for the adaptive value of social plasticity is to estimate the covariance between an individual’s plasticity in social behaviour in response to some environment factor and a fitness component of that individual. This approach uses multivariate mixed-effect modelling techniques (Arnold et al., 2019; Houslay & Wilson, 2017), which specifically account for repeated measures of the behaviour but allow a single measure of the fitness component *and* estimate the covariance at the correct hierarchical level (the among-individual level). Such models estimate the covariances between fitness and both individual trait means (via individual intercepts) and trait plasticities (via individual slopes), testing the adaptive benefit of social behaviours overall (i.e., if being more gregarious brings fitness benefits) and the adaptive benefit of social plasticity itself (i.e., if being more flexible in connectedness brings fitness benefits). This approach has been used to show the benefits of plasticity in traits such as life-history and phenology, for instance female Ural owls *Strix uralensis* that show steeper declines in clutch size with earlier lay date have higher fitness (Brommer et al., 2012). However, this approach has not yet been used for social behaviours. We therefore have no current evidence testing the suggestion that animals have evolved to plasticly adjust their social behaviour to buffer against environmental stressors.

Here, we test the associations between a fitness component (lifetime calving success) and both mean and plasticity of individual social behaviours, in wild female bottlenose dolphins (*Tursiops truncatus*). Bottlenose dolphins (*Tursiops* spp.) exhibit a fission-fusion grouping pattern where social interactions and relationships are critical components in their lives (Connor et al., 2000, 2019; Lusseau et al., 2006; Shane et al., 1986). Further, we have shown that dolphin gregariousness, quantified as the social network measure “strength”, increases in months of high salmon abundance (Fisher & Cheney, 2023, see also the correction: 2025), a key food source in this study population (Santos et al., 2001). This raises the question of whether this plasticity in response to salmon abundance is adaptive, i.e., does changing social behaviour enhance fitness. This would be expected if social associations influence the ability to acquire food (e.g., by cooperation or competition). Previous work has shown mean levels of social behaviour are associated with fitness components in toothed whales (Connor et al., 2022; Ellis et al., 2017; Gerber et al., 2022; Holmes et al., 2024) but the fitness benefits of plasticity itself are not quantified.

We directly tested for a possible adaptive function of social plasticity by fitting models that simultaneously estimate individual plasticity in social behaviours in response to salmon abundance and individual variance in a fitness component (calving success). Crucially, we estimate the covariance between calving success and both mean individual social behaviour (testing for benefits of mean behaviour) and plasticity in individual social behaviour in response to salmon abundance (testing for benefits of plasticity in behaviour). Since we previously looked at plasticity at both the monthly and yearly temporal scale (Fisher & Cheney, 2023), we repeat that two-scale approach to test whether responding to changes in monthly or yearly salmon abundance brings fitness benefits. Our aims are:

1. Determine if there are consequences for reproduction of changes in yearly salmon abundance.
2. Estimate plasticity in social behaviour in response to salmon abundance (as achieved by Fisher & Cheney, 2023 but with an updated dataset and with a slightly different modelling approach).
3. Estimate the association with fitness of individual variation in mean social behaviours.
4. Estimate the association with fitness of individual variation in plasticity in social behaviours.

## Methods

### Data collection

We used data from a long-term individual-based study of a population of over 220 bottlenose dolphins in the North Sea (Cheney et al., 2013, 2014, 2024). Boat-based photo-identification surveys have been carried out each year from May to September 1990 to 2024 within the Moray Firth Special Area of Conservation (SAC; 92/43/EEC). Upon encountering dolphins (an “encounter”), each group is carefully followed and photographs taken of both sides of as many of the dolphins’ dorsal fins as possible. Due to unique patterns of markings, scratches, and notches on each dolphin’s back and dorsal fin, individual dolphins are identified in these photographs and tracked through time. For this study, we used data from 13,259 sightings of individual dolphins across 2,492 encounters (average 70 groups per year, range 24 – 145, average group size 6, range 1 - 56).

Identifying individuals allows us to determine which dolphins are still alive, who is in the same group and so associating, and to identify new calves (Cheney et al., 2019). The year of birth and so age can be estimated for calves ≤ 2 years old, but ages are unknown for individuals first sighted as older calves, immature or adults. Mothers are assigned to calves based on repeat observations of the calf surfacing in echelon position next to the same adult female (Cheney et al., 2019), but paternity cannot be assigned as genetic data are not available. For more information on survey methods, see Cheney et al. (2014), while Fisher and Cheney (2023) provide information on using these survey records to construct social networks. Photographic survey work was conducted under NatureScot Animal Scientific Licences (latest number 228472) and all work was approved by the University of Aberdeen School of Biological Sciences ethics board (project ID SBSEC/01/2023/March).

### Social trait and fitness component calculation

We defined reproductive output per year in a binary manner as whether the female was known to give birth or not (Cheney et al., 2019). Sightings records are then used to determine whether the calf survived beyond one or two years. Calves are regularly observed with their mother in at least their first two years (Grellier et al., 2003) so repeat sightings of the mother without the calf at that age is a reliable indicator of calf death rather than dispersal. We estimated a fitness component “calving success”, a lifetime measure of the number of calves that survived more than two years produced by each female divided by the number of years she lived as a reproductive individual (year last observed – year of first reproduction). For females of known ages, the start of their reproductive period was eight years old (median age of first reproduction in this population) or the age when they had their first calf, whichever was younger; while for individuals of unknown age, it is from the year they were first observed with a calf or, if they never had a calf, eight years after they were first observed.

We can measure social traits of individuals by using the repeated observations of individuals in groups to build social networks. In a social network, individuals who are observed in the same group (photographed during an encounter, within 100m, engaged in similar activities and if travelling moving in a single direction; Wells et al., 1987) are connected, while those who are never observed in the same group are not connected (the “gambit of the group”; Whitehead & Dufault, 1999). The relative frequency two individuals are seen together defines the strength of the association – precisely, the number of times two individuals were seen together in a given time period divided by the total number of times both individuals were seen, either together or alone (the “simple ratio index”), which ranges from 0 to 1 (Cairns & Schwager, 1987). We constructed a social network per month and per year, using observations of individuals of either unknown age or above three years old, i.e., excluding known calves. We used the whole network to calculate two measures of each individual’s social network position, representing two dimensions of social behaviour. However, we then excluded individuals observed fewer than three times in that month or year from any analysis, as their social network position would be based on very few observations, and we excluded entire months (i.e., all rows in that month) if there were fewer than five groups observed in that month.

For each individual in each month or year, we calculated “strength”, the sum of all an individual’s weighted connections, giving an estimate of overall gregariousness. We also calculated “closeness”, which is based on the path lengths from an individual to all others in the network, weighted by the inverse of association strengths, representing connectedness to the entire population and has high values if an individual has strong connections linking them to all parts of the network (Opsahl et al., 2010). Both measures are therefore undirected (the association between individual A and B is identical to the association between B and A) and weighted (accounting for the intensity of associations in some way, rather than just presence/absence). We use these measures as we have previously found they show some variation in individual change in response to measures of salmon abundance, which is necessary to then be linked to among-individual differences in fitness (Fisher & Cheney, 2023). Fisher & Cheney (2023) also found that dolphin social behaviour varies with salmon abundance only at the monthly scale, and therefore we expect any association between calving success and plasticity to be at this scale. We plot social networks in Fig. 1 for 2011 and 2021 as examples.

**Figure 1.**
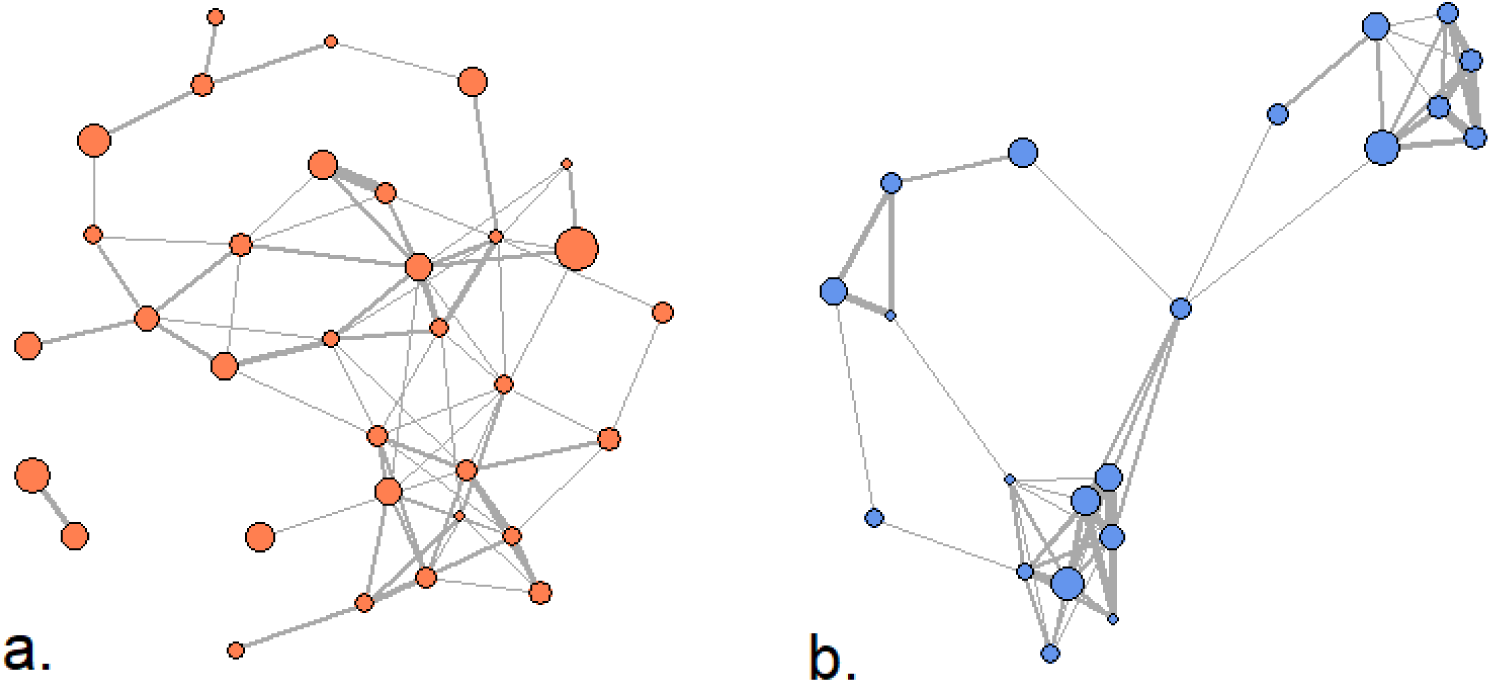
Plots of yearly social networks for 2011 (a.) and 2021 (b.). Points represent individual female dolphins and lines show weighted associations between them. We have only plotted associations with a simple ratio index above 0.2 to reduce visual clutter. Line thickness is increased with stronger associations, and point size is increased if that individual had higher lifetime calving success.

### Salmon abundance estimate

Salmon abundance in our study area (indeed the North Atlantic as a whole) has been undergoing a decline of around 70% in the last 25 years (ICES, 2017), but there is still considerable variation year to year (see Fig. S4 of Fisher & Cheney, 2023). We followed Fisher and Cheney (2023) and Lusseau *et al*. (2004) in using data on monthly catches by fishing rods (excluding nets) of adult Atlantic salmon (*Salmo salar*) from five rivers feeding into the Moray Firth (the Alness, Beauly, Canon, Ness, and Nairn) to quantify relative salmon abundance. Catch numbers peak in July-October (see Fig. S4 of Fisher & Cheney, 2023). These catch data are positively correlated among months and among rivers and with other data sources, suggesting they accurately reflect abundance (Thorley et al., 2005; Youngson et al., 2002). Further, salmon abundance in the current year is positively associated with group size (Lusseau et al., 2004) and in the current month with individual dolphin gregariousness (Fisher & Cheney, 2023). We downloaded the data from https://marine.gov.scot/data/marine-scotland-salmon-and-sea-trout-catches-salmon-district-shinyapp, using “rod data”, both one sea winter and multiple sea winter adults and summing retained and released fish for a monthly index of salmon abundance. We summed monthly catches within a calendar year for an annual index of salmon abundance.

### Data analysis

All models were fitted in MCMCglmm (v. 2.36) (Hadfield, 2010) in R (v. 4.4.1) (R Development Core Team & Team, 2016). To test the effect of salmon abundance on female reproduction (aim 1), we regressed the binary outcome of whether a female had a calf or not in that year on two variables: salmon abundance in the current year and salmon abundance in the previous year, as gestation in bottlenose dolphins is around 12 months and so could be affected by food availability in either calendar year. Both salmon abundances were z-transformed to a mean of zero and a standard deviation of one. In this analysis, we only included females who had been sighted a minimum of three times in that year, as we would otherwise be uncertain of their reproductive success. Note that for this and all other analyses increasing the minimum number of sightings per year decreases the number of observations and the number of unique females included. For the analysis of salmon abundance and reproductive success, if we require a minimum of three encounters per individual per year, we have 751 observations of 100 unique females, with four per year we have 646 observations of 95 unique females, and with five sightings per year we have 561 observations of 83 unique females. There is therefore an unavoidable trade-off between different aspects of statistical power. For this analysis the results were qualitatively the same with a threshold of three, four, or five sightings (Tables S1-3).

In our statistical models we included random effects of maternal ID and year to account for repeated observations of the same dolphins across years and for years that might have had higher or lower reproductive success on average. We also included a binary fixed effect of whether the female had given birth in the last two years or not, as this would greatly reduce the chances of her reproducing in the current year and needs to be accounted for (Cheney et al., 2019). We used a categorical error structure, and uninformative priors with V = 1 and nu = 0.02 for both random effects and the residual variance fixed at 1 (as in a model with a binary response the residual variance is defined by the mean). We ran the model for 110,000 iterations, discarding the first 10,000 as the model would initially be far from convergence, and using 1 out of every 100 remaining iterations for the posterior distribution to minimise autocorrelation. We checked model trace plots for satisfactory convergence and tested three runs of this model with the Gelman and Rubin convergence diagnostic. All terms achieved satisfactory convergence (potential scale reduction factor [PSRF] very close to 1). We report the posterior distribution medians in the main text as they are likely the least biased (Pick et al., 2023), but include the modes and means in tables reporting full model results in the supplementary materials.

We fitted four models to estimate mean and plasticity in social behaviour and testing for associations with calving success; aims 2-4, for all combinations of strength and closeness at monthly and yearly scales. All four models have an identical structure except that the monthly models also include a random effect of month, to account for seasonal variation in factors, aside from salmon abundance, that might influence social behaviour. For each model, we include a social network trait (either strength or closeness) and calving success as the two response variables. As before, we only included measures of social network traits where the female had been observed at least three times in a given month or year. Increasing this threshold to four or five approximately halves or quarters the dataset respectively for the monthly analyses, in theory making it harder to reliably estimate the parameters we are interested in (although in this case there was no impact as parameters were already hard to estimate, see Results and our power analysis). The effect is less dramatic in the yearly analysis as relatively more females have at least four or five sightings per year, but we keep the same threshold for both temporal scales to allow comparisons of the results. We also only included females in these analyses that had been seen for at least five years as reproductive, as some individuals below that threshold had artificially high calving success due to having a surviving calf in one year and then not being seen again.

We used salmon abundance in that year or month (z-transformed) as a fixed effect for the social network trait only. We include the random effects of individual ID for both traits and year (and month for the monthly models) for the social network trait only. We include random intercepts (which represent individual mean behaviour) for the social network trait and for calving success, and random slopes (which represent individual change in behaviour [plasticity]) for the social network trait with salmon abundance. We estimate all pairwise covariances between individual mean social behaviour, plasticity in individual social behaviour, and individual calving success. These are estimated at the among-individual level, and so the covariance between plasticity and calving success specifically tests whether individual flexibility is associated with fitness (Houslay & Wilson, 2017). We used Gaussian error structures for both traits, uninformative priors with V = 1 and nu = 0.02 for the residual variance of the social network trait and the random effect of year. The residual variance for calving success was fixed at 0.0001 to aid estimation as the trait does not vary within individuals. Finally, for the variance-covariance matrix at the among-individual level, we used a parameter-expanded prior with V = 1, nu = 3, alpha.mu = 0, and alpha.V = 625 following Houslay and Wilson (2017). We checked the trace plots for all four models for satisfactory convergence and tested three runs of each model with the Gelman and Rubin convergence diagnostic. Terms in all models achieved satisfactory convergence (PSRF < 1.04), apart from the random effect of month in the model for monthly closeness (PSRF = 1.08). Since this is a control term and not of specific interest we do not believe this is problematic.

### Power analysis

After viewing our results results, we decided to conduct a power analysis to assess our ability to detect correlations of a reasonable size. We did this based on Pick et al. (2023), and following discussions with J. Pick. We aimed to calculate the minimum detectable correlations (MDCs) – the smallest correlations (either side of zero) our dataset could reliably detect. These would be beyond the 95% credible intervals of the null distributions for each correlation of interest. The null distributions need to be estimated based on a randomised dataset, featuring the same characteristics as the original dataset, but containing a covariance between calving success and random slope intercepts or slopes that was genuinely 0. To create these, we randomly permuted calving success values among individuals within our datasets (separately for both yearly and monthly scales) but kept all other features the same. This step breaks any possible association between mean or plasticity in social behaviour and calving success.

We then fitted the same models as in our main analysis and extracted a single random value from each of the posterior distributions for the covariances between individual calving success and both mean behaviour and plasticity. We repeated this 1000 times, giving 1000 estimates for the covariance between calving success and mean social behaviour and the covariance between calving success and plasticity in social behaviour from datasets where there are no relationships. We repeated this once each for strength and closeness at both the monthly and yearly scales, giving eight sets of covariances in total. The spread of each distribution indicates our power to detect non-zero covariances – a narrow spread indicates good power and a wide spread low power. We convert these to correlations to aid interpretation by dividing by the square root of the product of the estimates of the variance in individual mean behaviour (intercepts) and individual plasticities (slopes) from the original dataset (reported in Tables S8-11). The MDCs are those below or above the 2.5% and 97.5% percentiles of the null distribution respectively.

## Results

To test the effect of salmon abundance on yearly calf production and survival, we had 751 observations of known reproductive status across 100 different females. Salmon abundance in both the current year (posterior distribution median [PDM] = - 0.153, 95% credible intervals [CIs] = -0.599 to 0.241) and previous year (PDM = - 0.133, 95% CIs = -0.614 to 0.306) did not influence reproductive success. Having a calf in the previous two years reduced the chance of a female producing a calf (PDM = -2.131, 95% CIs = -2.813 to -1.470; full model results in Table S1).

To test for the fitness associations between mean and plasticity in social behaviour at the monthly scale, we had 711 measures of social behaviour across 62 different females. For the yearly scale analysis, we had 619 measures of social behaviour across 65 different females.

At the monthly level, there was no evidence that individual differences in calving success were associated with individual differences in plasticity or mean behaviour for strength or closeness (Figs. 2a & b & Table 1). There were also no associations between individual mean behaviour and plasticity of behaviour for either trait (Table 1). Full model results are given in Tables S4 & 5.

**Figure 2.**
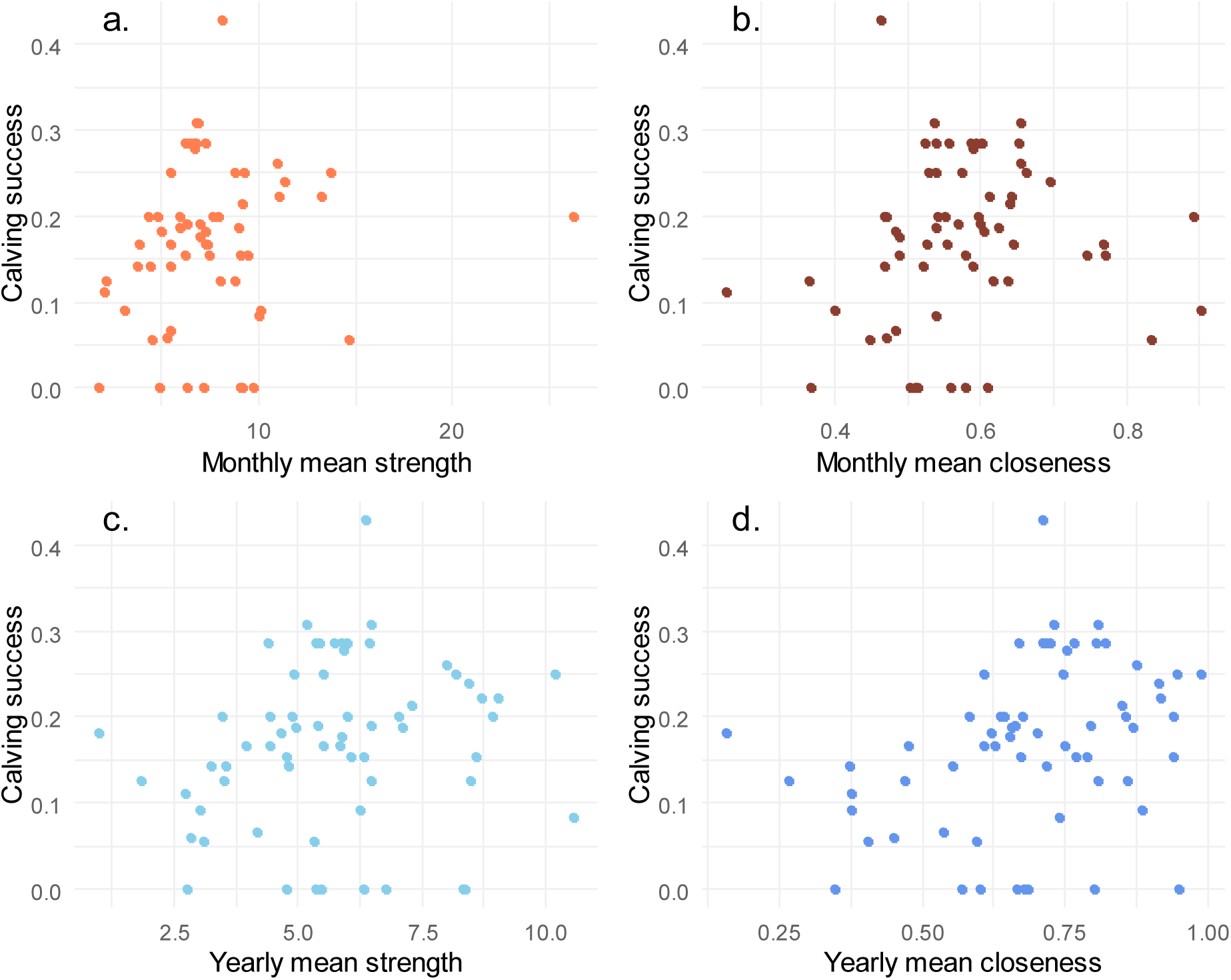
Plots of the relationships between monthly (a. & b.) and yearly (c. & d.) means of social network traits (a. & c strength, b. & d. closeness) and calving success. Each point represents an individual females’ mean social network trait for a given month or year against that individual’s calving success (one lifetime measure per individual) (plotted in ggplot2; Wickham, 2009).

**Table 1.** Estimates of the covariance parameters for the models with strength and closeness. In each cell are the posterior distribution median (and the 95% credible intervals in parentheses) for the covariance between the two terms shown. The diagonal is filled with Xs, above it are values for the monthly analyses, while below are values for the yearly analyses.

|  | *Strength* |  |  |
| --- | --- | --- | --- |
|  | **Individual mean behaviour** | **Plasticity in individual behaviour** | **Calving success** |
| **Individual mean behaviour** | X | -0.120 (-0.878 to 0.268) | 0.028 (-0.027 to 0.086) |
| **Plasticity in individual behaviour** | -0.020 (-0.247 to 0.106) | X | -0.005 (-0.035 to 0.013) |
| **Calving success** | 0.031 (-0.006 to 0.076) | -0.000 (-0.016 to 0.012) | X |
|  | *Closeness* |  |  |
|  | **Individual mean behaviour** | **Plasticity in individual behaviour** | **Calving success** |
| **Individual mean behaviour** | X | -0.000 (-0.001 to 0.000) | 0.000 (-0.001 to 0.003) |
| **Plasticity in individual behaviour** | -0.001 (-0.002 to 0.000) | X | -0.000 (-0.002 to 0.001) |
| **Calving success** | 0.002 (-0.000 to 0.005) | -0.001 (-0.002 to 0.001) | X |

At the yearly level, there were no associations between individual differences in calving success and individual differences in plasticity or mean behaviour for strength and closeness (Table 1). There were also no covariances between individuals’ mean behaviour and plasticity of behaviour for either trait (Table 1). Full model results are given in Tables S6 & 7.

The power analysis revealed that the MDCs for the relationship between calving success and mean strength at the monthly scale were -0.569 and 0.591 i.e., we could reliably detect correlations lower than -0.569 or above 0.591. For calving success and plasticity in strength at the same temporal scale the MDCs were -1.023 and 1.022 i.e., beyond -1 and 1 – essentially undetectable. The MDCs for closeness at the monthly scale were slightly larger (mean closeness MDCs -0.693 and 0.709, plasticity in closeness MDCs -1.169 and 1.047). We saw slightly lower MDCs for correlations between calving success and mean behaviour at the yearly scale (strength MDCs -0.473 and 0.439, closeness MDCs -0.467 and 0.515). We saw very low power for the correlation between calving success and plasticity in strength (MDCs 1.500 and 1.713) but, in contrast to the monthly scale, some ability to detect very strong correlations with plasticity in closeness at the yearly scale (MDCs -0.878 and 0.946). Therefore, MDCs for the relationships between mean behaviour and calving success were around 0.5, indicating we could only detect effects above medium strength. Meanwhile, we had very little ability to detect relationships between plasticity in behaviour and calving success (3/4 sets of MDCs were beyond -1 to 1).

## Discussion

We found that salmon abundance was not associated with female reproductive success (aim 1). We estimated both individual means and plasticities of social behaviour (aim 2) and found the individual means were not associated with female calving success at either monthly or yearly timescales (aim 3). We also found no evidence that social plasticity might be adaptive as there were no covariances between plasticity in strength or closeness in response to salmon abundance and calving success (aim 4).

Female dolphins who had high strength and closeness over an entire year had similar calving success to those with low network metrics. This matches results from Stanton and Mann’s (2012) study on a population of Indo-Pacific bottlenose dolphins *T. aduncus*, where females showed a non-significant relationship between eigenvector centrality (a social network measure of connectedness accounting for the connections possessed by one’s social associates) and survival (in comparison to males, who showed a positive relationship). In addition, recent research on the same population showed that adult males with strong bonds who are well integrated in their cooperative alliance sire more offspring (Gerber et al., 2022). Given that we studied different fitness components and a different species with a different ecology to Stanton and Mann and Gerber et al., whether our results are directly comparable is in question. Further comparative work on associations between fitness components and social behaviour in both sexes and multiple populations of *Tursiops* spp. is warranted.

We did not detect any fitness benefits to plasticity. Female dolphins do not appear to adjust their social behaviour to buffer low salmon abundance or take advantage of high salmon abundance. This observational dataset has inherent limitations, for example, data was only available for the summer months, not all females are observed every year, not all calves are identified, and some calves may die before they are observed. These limitations add noise to the dataset and may prevent us detecting any genuine relationships between this fitness component and other traits. Additionally, given we show that salmon abundance is not related to calf production, it is perhaps not surprising that changes in social behaviour are not required to “buffer” low salmon numbers. Dolphins may switch to other prey when salmon are scarce, such as whiting *Merlangius merlangus*, saithe *Pollachius virens*, and cod *Gadus morhua* (Santos et al., 2001), instead of changing their social behaviour. Estimating the resource requirements of the population should be a priority in the face of a changing climate. Meanwhile, social behaviour, which shows considerable within-individual variation, may instead change in response to reproductive opportunities, to shifts in local currents and fronts, or other environmental factors such as anthropogenic disturbance, and the adaptive value of plasticity might show itself in any of these contexts.

The power analysis reveals that, even in a long-term study with an extensive dataset, filtered to only include individuals with reliable observations, we would struggle to detect social buffering (the correlation between individual plasticity and calving success) even if it was there. There was more power to estimate associations between mean social behaviour and calving success, yet we still found no relationships. In contrast, positive associations between mean social behaviour and fitness components, such as longevity, have been reported multiple times in terrestrial mammals, such as rhesus macaques *Macaca mulatta* (Ellis et al., 2019), big horn sheep *Ovis canadensis* (Wal et al., 2015) and spotted hyenas *Crocuta crocuta* (Yoshida et al., 2016). In general, fitness benefits of sociality are thought to arise due to detection of and protection from predators, opportunities to share information, and the ability to complete tasks more efficiently compared to when alone (Rubenstein & Abbot, 2017). These factors may be absent in our population. Social grouping is unlikely to function as protection from predators, as their natural predators are absent from these waters (Cheney et al., 2019). Additionally, while the dolphins in the Moray Firth SAC tend to be observed in groups (median size approximately 6-10 individuals; Cheney et al., 2024), they are often observed hunting individually (B. Cheney, pers. obs.), suggesting grouping enhancing hunting efficiency may not be essential.

As far as we are aware, our study is the first to estimate the covariance between plasticity in social behaviour and a fitness component. Our finding of (very) low power is a warning to other studies looking to estimate this relationship. The only study systems that might reliably detect social buffering (in this conceptualisation) would have exceptionally consistent differences among individuals in plasticity of social behaviour (to reliably estimate individual slopes), reliable estimates of fitness components, and a strong relationship between the two. This may well be hard to achieve, and so alternative options to explore equivalent patterns should be considered. Two studies on birds have tested whether different social behaviours yield fitness benefits in different conditions. In superb starlings *Lamprotornis superbus*, males have higher survival in larger groups only in wet years but the reverse is true in dry years (Guindre-Parker & Rubenstein, 2020), while in females, larger groups are always associated with higher survival. Meanwhile, in opposition to predictions that social behaviours help buffer harsh conditions, Seychelles warblers *Acrocephalus sechellensis* show lower rates of reproduction with low rainfall, but in years of little rain alloparenting was *less* common (Borger et al., 2023). However, as previously mentioned, these type of analyses do not explicitly test whether plasticity itself is associated with fitness. Further, the results are not consistent with the idea of increasing social behaviour universally bringing fitness benefits in harsher conditions. While there is a range of evidence to show that social behaviour plastically varies with environmental conditions (Fisher et al., 2021), there is currently no evidence that this plasticity buffers harsh environmental conditions.

In general, there is variable evidence for selection on plasticity. For example, Pelletier *et al*. (2007) found selection on seasonal plasticity in female bighorn sheep ewes, with more plastic individuals having higher reproduction. Meanwhile, female red deer *Cervus elaphus* that give birth early after dry autumns and late after wet autumns may have higher fitness, but this trend was marginally non-significant (p = 0.06) and only applied if those females were born in a high density period (Nussey et al., 2005). Meanwhile, female Ural owls show selection on quadratic reaction norms between laying date and clutch size, but no selection on the linear reaction norms (Brommer et al., 2012), which is what we evaluated here. These studies give an idea of the diversity of findings, but we are not aware of any meta-analyses collecting selection estimates on plasticity in animals. Arnold et al. (2019) reviewed evidence for selection on thermal plasticity in plants and found it to be lacking, while Davidson et al. (2011) found that invasive plants were more plastic than non-invasive plants, but that this plasticity did not confer especial fitness benefits. Therefore, while we have spent much of the 21st century wondering what the costs of plasticity are (Pigliucci, 2005), the benefits might not be as empirically well-supported as expected.

Future studies on the benefits of social plasticity in bottlenose dolphins could focus on “developmental plasticity” – the ability of a genotype to produce different phenotypes over the course of development, which cannot be reversed (Taborsky, 2017). This is in contrast to “reversible plasticity”, where an adult individual changes its phenotype and can reverse the changes back (measured here). For example, stress exposure in juvenile rats *Rattus norvegicus* reduces sociality in later life (Toth et al., 2008). Evidence that developmental plasticity in social behaviour is adaptive, however, is inconclusive, with seemingly maladaptive changes often observed (Spencer, 2017). In Indo-Pacific bottlenose dolphins, juveniles can show considerable variation in social behaviour (Gibson & Mann, 2008), which persists into adulthood (Evans et al., 2021), and has fitness consequences (Gerber et al., 2022; Holmes et al., 2024; Stanton & Mann, 2012). Further work should therefore identify what factors drive early variation in social behaviour and the adaptive consequences of early-life effects.

## Acknowledgements

We are forever grateful to Paul Thompson for conception and development of this long-term individual-based research study, and for commenting on this manuscript. We are especially thankful for our colleagues at the Lighthouse Field Station, past and present, who have contributed to and advanced this long-term study. Simon Allen and four anonymous reviewers made numerous useful comments and suggestions on an earlier draft. The contribution of numerous funders has been key for this long-term study, including University of Aberdeen, NatureScot, Beatrice Offshore Windfarm Ltd., Moray Offshore Renewables Ltd., Marine Scotland, the Crown Estate, Highlands and Islands Enterprise, the BES, ASAB, Greenpeace Environmental Trust, Whale and Dolphin Conservation, Talisman Energy (UK) Ltd., Department of Energy and Climate Change, Chevron and Natural Environment Research. Survey work was conducted under NatureScot Animal Scientific Licences.

## Authors’ contributions

BJC collects data on this long-term project and curates the database. DNF conceived the idea of this particular manuscript and conducted preliminary analyses. DNF and BJC discussed the ideas and the analysis before DNF finalised the analysis and wrote a draft. BJC edited the draft and together the authors revised the final version.

## References

Arnold, P. A., Nicotra, A. B., & Kruuk, L. E. B. (2019). Sparse evidence for selection on phenotypic plasticity in response to temperature. Philosophical Transactions of the Royal Society B: Biological Sciences, 374(1768), 20180185. 10.1098/rstb.2018.0185

Auer, S. K., Salin, K., Rudolf, A. M., Anderson, G. J., & Metcalfe, N. B. (2015). Flexibility in metabolic rate confers a growth advantage under changing food availability. Journal of Animal Ecology, 84(5), 1405–1411. 10.1111/1365-2656.12384

Blumstein, D. T., Hayes, L. D., & Pinter-Wollman, N. (2023). Social consequences of rapid environmental change. Trends in Ecology & Evolution, 38(4), 337–345. 10.1016/j.tree.2022.11.005

Borger, M. J., Richardson, D. S., Dugdale, H., Burke, T., & Komdeur, J. (2023). Testing the environmental buffering hypothesis of cooperative breeding in the Seychelles warbler. Acta Ethologica, 26(3), 211–224. 10.1007/s10211-022-00408-y

Brommer, J. E., Kontiainen, P., & Pietiäinen, H. (2012). Selection on plasticity of seasonal life-history traits using random regression mixed model analysis. Ecology and Evolution, 2(4), 695–704. 10.1002/ece3.60

Cairns, S. J., & Schwager, S. J. (1987). A comparison of association indices. Animal Behaviour, 35(5), 1454–1469. 10.1016/S0003-3472(87)80018-0

Candolin, U., & Wong, B. B. M. (2011). Behavioural Responses to a Changing World: Mechanisms & Consequences (U. Candolin & B. B. M. Wong, Eds.). Oxford University Press. https://global.oup.com/academic/product/behavioural-responses-to-a-changing-world-9780199602575?cc=gb&lang=en&

Cheney, B. J., Arso Civil, M., Hammond, P. S., & Thompson, P. M. (2024). Site Condition Monitoring of bottlenose dolphins within the Moray Firth Special Area of Conservation 2017-2022 (No. 1360; NatureScot Research Report). NatureScot. https://www.nature.scot/doc/naturescot-research-report-1360-site-condition-monitoring-bottlenose-dolphins-within-moray-firth

Cheney, B. J., Corkrey, R., Durban, J. W., Grellier, K., Hammond, P. S., Islas-Villanueva, V., Janik, V. M., Lusseau, S. M., Parsons, K. M., Quick, N. J., Wilson, B., & Thompson, P. M. (2014). Long-term trends in the use of a protected area by small cetaceans in relation to changes in population status. Global Ecology and Conservation, 2, 118–128. 10.1016/j.gecco.2014.08.010

Cheney, B. J., Thompson, P. M., & Cordes, L. S. (2019). Increasing trends in fecundity and calf survival of bottlenose dolphins in a marine protected area. Scientific Reports, 9(1), 1767. 10.1038/s41598-018-38278-9

Cheney, B. J., Thompson, P. M., Ingram, S. N., Hammond, P. S., Stevick, P. T., Durban, J. W., Culloch, R. M., Elwen, S. H., Mandleberg, L., Janik, V. M., Quick, N. J., Islas-Villanueva, V., Robinson, K. P., Costa, M., Eisfeld, S. M., Walters, A., Phillips, C., Weir, C. R., Evans, P. G. H., … Wilson, B. (2013). Integrating multiple data sources to assess the distribution and abundance of bottlenose dolphins Tursiops truncatus in Scottish waters. Mammal Review, 43(1), 71–88. 10.1111/j.1365-2907.2011.00208.x

Connor, R. C., Krützen, M., Allen, S. J., Sherwin, W. B., & King, S. L. (2022). Strategic intergroup alliances increase access to a contested resource in male bottlenose dolphins. Proceedings of the National Academy of Sciences, 119(36), e2121723119. 10.1073/pnas.2121723119

Connor, R. C., Sakai, M., Morisaka, T., & Allen, S. J. (2019). The Indo-Pacific Bottlenose Dolphin (Tursiops aduncus). In B. Würsig (Ed.), Ethology and Behavioral Ecology of Odontocetes (pp. 345–368). Springer International Publishing. 10.1007/978-3-030-16663-2_16

Connor, R. C., Wells, R. S., Mann, J., & Read, A. J. (2000). The bottlenose dolphin. In Cetacean Societies: Field studies of whales and dolphins (pp. 91–125). University of Chicago Press Chicago, Illinois. https://books.google.com/books?hl=en&lr=&id=W-UQNoxMONwC&oi=fnd&pg=PA91&dq=The+bottlenose+dolphin.+In%C2%A0Cetacean+Societies:+Field+studies+of+whales+and+dolphins&ots=WVoTXSwbyk&sig=ElaUH2Qk9T9v3N_aTx_-csPPcpc

Davidson, A. M., Jennions, M., & Nicotra, A. B. (2011). Do invasive species show higher phenotypic plasticity than native species and, if so, is it adaptive? A meta-analysis. Ecology Letters, 14(4), 419–431. 10.1111/j.1461-0248.2011.01596.x

Ellis, S., Franks, D. W., Nattrass, S., Cant, M. A., Weiss, M. N., Giles, D., Balcomb, K. C., & Croft, D. P. (2017). Mortality risk and social network position in resident killer whales: Sex differences and the importance of resource abundance. *Proceedings*. Biological Sciences, 284(1865), 20171313. 10.1098/rspb.2017.1313

Ellis, S., Snyder-Mackler, N., Ruiz-Lambides, A., Platt, M. L., & Brent, L. J. N. (2019). Deconstructing sociality: The types of social connections that predict longevity in a group-living primate. Proceedings of the Royal Society B: Biological Sciences, 286(1917), 20191991. 10.1098/rspb.2019.1991

Evans, T., Krzyszczyk, E., Frère, C., & Mann, J. (2021). Lifetime stability of social traits in bottlenose dolphins. Communications Biology, 4(1), 1–8. 10.1038/s42003-021-02292-x

Firman, R. C., Rubenstein, D. R., Moran, J. M., Rowe, K. C., & Buzatto, B. A. (2020). Extreme and Variable Climatic Conditions Drive the Evolution of Sociality in Australian Rodents. Current Biology, 30(4), 691–697.e3. 10.1016/j.cub.2019.12.012

Fisher, D. N., & Cheney, B. J. (2023). Dolphin social phenotypes vary in response to food availability but not the North Atlantic Oscillation index. Proceedings of the Royal Society B: Biological Sciences, 290(2008), 20231187. 10.1098/rspb.2023.1187

Fisher, D. N., & Cheney, B. J. (2025). Correction to: ‘Dolphin social phenotypes vary in response to food availability but not the North Atlantic Oscillation index’ (2023), by Fisher and Cheney. Proceedings of the Royal Society B: Biological Sciences, 292(2046), 20242078. 10.1098/rspb.2024.2078

Fisher, D. N., Kilgour, R. J., Siracusa, E. R., Foote, J. R., Hobson, E. A., Montiglio, P. O., Saltz, J. B., Wey, T. W., & Wice, E. W. (2021). Anticipated effects of abiotic environmental change on intraspecific social interactions. Biological Reviews, 96(6), 2661–2693. 10.1111/brv.12772

Gerber, L., Connor, R. C., Allen, S. J., Horlacher, K., King, S. L., Sherwin, W. B., Willems, E. P., Wittwer, S., & Krützen, M. (2022). Social integration influences fitness in allied male dolphins. Current Biology, 32(7), 1664–1669.

Gibson, Q. A., & Mann, J. (2008). Early social development in wild bottlenose dolphins: Sex differences, individual variation and maternal influence. Animal Behaviour, 76(2), 375–387. 10.1016/j.anbehav.2008.01.021

Grellier, K., Hammond, P. S., Wilson, B., Sanders-Reed, C. A., & Thompson, P. M. (2003). Use of photo-identification data to quantify mother–calf association patterns in bottlenose dolphins. Canadian Journal of Zoology, 81(8), 1421–1427. 10.1139/z03-132

Guevara, J., & Avilés, L. (2015). Ecological predictors of spider sociality in the Americas. Global Ecology and Biogeography, 24(10), 1181–1191. 10.1111/geb.12342

Guindre-Parker, S., & Rubenstein, D. R. (2020). Survival Benefits of Group Living in a Fluctuating Environment. The American Naturalist, 195(6), 1027–1036. 10.1086/708496

Hadfield, J. D. (2010). MCMC methods for multi-response generalized linear mixed models: The MCMCglmm R package. Journal of Statistical Software, 33(2), 1–22.

Hogstad, O. (1984). Variation in numbers, territoriality and flock size of a Goldcrest Regulus regulus population in winter. Ibis, 126(3), 296–306. 10.1111/j.1474-919X.1984.tb00252.x

Holmes, K. G., Krützen, M., Ridley, A. R., Allen, S. J., Connor, R. C., Gerber, L., Flaherty Stamm, C., & King, S. L. (2024). Juvenile social play predicts adult reproductive success in male bottlenose dolphins. Proceedings of the National Academy of Sciences, 121(25), e2305948121. 10.1073/pnas.2305948121

Houslay, T. M., & Wilson, A. J. (2017). Avoiding the misuse of BLUP in behavioural ecology. Behavioral Ecology, 12(Pt A), S9. 10.1093/beheco/arx023

ICES. (2017). Report of the working group on North Atlantic salmon (WGNAS). In ICES CM/ACOM*. vol. 20* (p. 296). ICES International Council for the Exploration of the Sea Copenhagen, Denmark.

Jetz, W., & Rubenstein, D. R. (2011). Environmental Uncertainty and the Global Biogeography of Cooperative Breeding in Birds. Current Biology, 21(1), 72–78. 10.1016/J.CUB.2010.11.075

Komdeur, J., & Ma, L. (2021). Keeping up with environmental change: The importance of sociality. Ethology, 127(10), 790–807. 10.1111/ETH.13200

López-Sepulcre, A., & Kokko, H. (2012). Understanding behavioural responses and their consequences. In U. Candolin & B. B. M. Wong (Eds.), Behavioural Responses to a Changing World: Mechanisms and Consequences. Oxford University Press. https://www.oxfordscholarship.com/view/10.1093/acprof:osobl/9780199602568.001.0001/acprof-9780199602568-chapter-1

Lusseau, D., Williams, R., Wilson, B., Grellier, K., Barton, T. R., Hammond, P. S., & Thompson, P. M. (2004). Parallel influence of climate on the behaviour of Pacific killer whales and Atlantic bottlenose dolphins. Ecology Letters, 7(11), 1068–1076. 10.1111/j.1461-0248.2004.00669.x

Lusseau, D., Wilson, B., Hammond, P. S., Grellier, K., Durban, J. W., Parsons, K. M., Barton, T. R., & Thompson, P. M. (2006). Quantifying the influence of sociality on population structure in bottlenose dolphins. Journal of Animal Ecology, 75(1), 14–24. 10.1111/j.1365-2656.2005.01013.x

Nussey, D. H., Clutton-Brock, T. H., Elston, D. A., Albon, S. D., & Kruuk, L. E. B. (2005). Phenotypic Plasticity in a Maternal Trait in Red Deer. Journal of Animal Ecology, 74(2), 387–396.

Opsahl, T., Agneessens, F., & Skvoretz, J. (2010). Node centrality in weighted networks: Generalizing degree and shortest paths. Social Networks, 32(3), 245–251. 10.1016/j.socnet.2010.03.006

Pelletier, F., Réale, D., Garant, D., Coltman, D. W., & Festa-Bianchet, M. (2007). Selection on heritable seasonal phenotypic plasticity of body mass. Evolution, 61(8), 1969–1979. 10.1111/j.1558-5646.2007.00160.x

Pick, J. L., Kasper, C., Allegue, H., Dingemanse, N. J., Dochtermann, N. A., Laskowski, K. L., Lima, M. R., Schielzeth, H., Westneat, D. F., Wright, J., & Araya-Ajoy, Y. G. (2023). Describing posterior distributions of variance components: Problems and the use of null distributions to aid interpretation. Methods in Ecology and Evolution, 14(10), 2557–2574. 10.1111/2041-210X.14200

Pigliucci, M. (2005). Evolution of phenotypic plasticity: Where are we going now? Trends in Ecology & Evolution, 20(9), 481–486.

R Development Core Team, & Team, R. D. C. (2016). R: A language and environment for statistical computing. In R Foundation for Statistical Computing (Vol. 1, Issue 3.1.3, p. 409) [Computer software]. R Foundation for Statistical Computing. 10.1007/978-3-540-74686-7

Rubenstein, D. R., & Abbot, P. (2017). Comparative Social Evolution (D. R. Rubenstein & P. Abbot, Eds.). Cambridge University Press. 10.1017/9781107338319

Santos, M. B., Pierce, G. J., Reid, R. J., Patterson, I. a. P., Ross, H. M., & Mente, E. (2001). Stomach contents of bottlenose dolphins (Tursiops truncatus) in Scottish waters. Journal of the Marine Biological Association of the United Kingdom, 81(5), 873–878. 10.1017/S0025315401004714

Shane, S. H., Wells, R. S., & Würsig, B. (1986). Ecology, Behavior and Social Organization of the Bottlenose Dolphin: A Review. Marine Mammal Science, 2(1), 34–63. 10.1111/j.1748-7692.1986.tb00026.x

Sheehan, M. J., Botero, C. A., Hendry, T. A., Sedio, B. E., Jandt, J. M., Weiner, S., Toth, A. L., & Tibbetts, E. A. (2015). Different axes of environmental variation explain the presence vs. Extent of cooperative nest founding associations in *Polistes* paper wasps. Ecology Letters, 18(10), 1057–1067. 10.1111/ele.12488

Sih, A. (2013). Understanding variation in behavioural responses to human-induced rapid environmental change: A conceptual overview. Animal Behaviour. https://www.sciencedirect.com/science/article/pii/S0003347213000936?casa_token=2-UuMOtyassAAAAA:eHXDj44BrAgcu7HfDrip6_IzwDas_b4SSR0q6EJbwryT_cas6MqS3IB_cBN33xRlfTVSKNOxmg

Sih, A., Ferrari, M. C. O., & Harris, D. J. (2011). Evolution and behavioural responses to human-induced rapid environmental change. Evolutionary Applications, 4(2), 367–387. 10.1111/j.1752-4571.2010.00166.x

Spencer, K. A. (2017). Developmental stress and social phenotypes: Integrating neuroendocrine, behavioural and evolutionary perspectives. Philosophical Transactions of the Royal Society B: Biological Sciences, 372(1727), 20160242. 10.1098/rstb.2016.0242

Stanton, M. A., & Mann, J. (2012). Early social networks predict survival in wild bottlenose dolphins. PloS One, 7(10), e47508. 10.1371/journal.pone.0047508

Székely, T. (Tamás), Moore, A. J. (Allen J., & Komdeur, J. (2010). Social behaviour: Genes, ecology and evolution. Cambridge University Press. https://www.cambridge.org/gb/academic/subjects/life-sciences/animal-behaviour/social-behaviour-genes-ecology-and-evolution?format=PB&isbn=9780521709620

Taborsky, B. (2017). Chapter Three - Developmental Plasticity: Preparing for Life in a Complex World. In M. Naguib, J. Podos, L. W. Simmons, L. Barrett, S. D. Healy, & M. Zuk (Eds.), Advances in the Study of Behavior (Vol. 49, pp. 49–99). Academic Press. 10.1016/bs.asb.2016.12.002

Thorley, J. L., Eatherley, D. M. R., Stephen, A. B., Simpson, I., MacLean, J. C., & Youngson, A. F. (2005). Congruence between automatic fish counter data and rod catches of Atlantic salmon (Salmo salar) in Scottish rivers. ICES Journal of Marine Science, 62(4), 808–817. 10.1016/j.icesjms.2005.01.016

Toth, E., Avital, A., Leshem, M., Richter-Levin, G., & Braun, K. (2008). Neonatal and juvenile stress induces changes in adult social behavior without affecting cognitive function. Behavioural Brain Research, 190(1), 135–139. 10.1016/j.bbr.2008.02.012

Tuomainen, U., & Candolin, U. (2010). Behavioural responses to human-induced environmental change. Biological Reviews, 86(3), 640–657. 10.1111/j.1469-185X.2010.00164.x

Wal, E. V., Festa-Bianchet, M., Vander Wal, E., Festa-Bianchet, M., Réale, D., Coltman, D. W., & Pelletier, F. (2015). Sex-based differences in the adaptive value of social behavior contrasted against morphology and environment. Ecology, 96(3), 631–641. 10.1890/14-1320.1

Warrington, M., Fisher, D., Komdeur, J., Pilakouta, N., & Griesser, M. (2024). Stronger together? A framework for studying population resilience to climate change impacts via social shielding. Life Sciences. 10.32942/X2QG9C

Wells, R. S., Scott, M. D., & Irvine, A. B. (1987). The Social Structure of Free-Ranging Bottlenose Dolphins. In H. H. Genoways (Ed.), Current Mammalogy (pp. 247–305). Springer US. 10.1007/978-1-4757-9909-5_7

Whitehead, H., & Dufault, S. (1999). Techniques for Analyzing Vertebrate Social Structure Using Identified Individuals: Review and Recommendations. Advances in the Study of Behavior, 28. http://www.sciencedirect.com/science/article/pii/S0065345408602156

Wickham, H. (2009). ggplot2: Elegant graphics for data analysis [Computer software]. Springer New York. http://had.co.nz/ggplot2/book

Wong, B. B. M., & Candolin, U. (2015). Behavioral responses to changing environments. Behavioral Ecology. 10.1093/beheco/aru183

Yoshida, K. C. S., Van Meter, P. E., & Holekamp, K. E. (2016). Variation among free-living spotted hyenas in three personality traits. Behaviour, 153(13–14), 1665–1722. 10.1163/1568539X-00003367

Youngson, A. F., MacLean, J. C., & Fryer, R. J. (2002). Rod catch trends for early-running MSW salmon in Scottish rivers (1952–1997): Divergence among stock components. ICES Journal of Marine Science, 59(4), 836–849. 10.1006/jmsc.2002.1195

