## Supplementary materials for "Examining the fitness benefits of social plasticity to prey availability in bottlenose dolphins"

There are seven supplementary tables to accompany the main text.

Table S1 gives the full model results for the analysis of the effect of salmon abundance on calf production (with a minimum threshold of 3 sightings per year) while Tables S2 & 3 give the same analysis with thresholds of 4 and 5 sightings per year respectively.

Tables S4 & 5 give the full model results for the monthly analysis of strength and closeness.

Tables S6 & 7 give the full model results for the yearly analysis of strength and closeness.

**Table S1.** Full model results of the analysis of salmon abundance in current year (“Current salmon”) and in the previous year (“Previous salmon) with a minimum sighting threshold of three per year (the analysis reported in the main text). For each effect we give the posterior distribution (PD) mode, median, and mean, the lower and upper 95% credible intervals (CI), and the effective sample size. For the fixed effects we also give the pMCMC values.

| Variable | PD mode | PD median | PD mean | Lower 95% CI | Upper 95% CI | Effective sample size | pMCMC | Effect type |
| --- | --- | --- | --- | --- | --- | --- | --- | --- |
| Intercept | -1.5300 | -1.4495 | -1.4581 | -1.8537 | -1.0766 | 765.1795 | 0.0010 | fixed |
| Current salmon | -0.2088 | -0.1532 | -0.1543 | -0.5986 | 0.2413 | 1000.0000 | 0.4700 | fixed |
| Previous salmon | -0.0743 | -0.1331 | -0.1427 | -0.6139 | 0.3057 | 1000.0000 | 0.5060 | fixed |
| If the female had a calf in the last two years | -2.0460 | -2.1312 | -2.1308 | -2.8128 | -1.4698 | 648.9902 | 0.0010 | fixed |
| Random effect mother ID | 0.0256 | 0.0959 | 0.1622 | 0.0021 | 0.5304 | 354.5740 | NA | random |
| Random effect year | 0.3879 | 0.4220 | 0.5066 | 0.0032 | 1.1831 | 573.8768 | NA | random |

**Table S2.** Full model results of the analysis of salmon abundance in current year (“Current salmon”) and in the previous year (“Previous salmon) with a minimum sighting threshold of four per year. For each effect we give the posterior distribution (PD) mode, median, and mean, the lower and upper 95% credible intervals (CI), and the effective sample size. For the fixed effects we also give the pMCMC values.

| Variable | PD mode | PD median | PD mean | Lower 95% CI | Upper 95% CI | Effective sample size | pMCMC | Effect type |
| --- | --- | --- | --- | --- | --- | --- | --- | --- |
| Intercept | -1.5300 | -1.4495 | -1.3652 | -1.7841 | -1.0211 | 587.8328 | 0.0010 | fixed |
| Current salmon | -0.2088 | -0.1532 | -0.2092 | -0.6679 | 0.1686 | 1000.0000 | 0.3180 | fixed |
| Previous salmon | -0.0743 | -0.1331 | -0.1167 | -0.5332 | 0.3056 | 1000.0000 | 0.6100 | fixed |
| If the female had a calf in the last two years | -2.0460 | -2.1312 | -2.2665 | -3.0383 | -1.5945 | 745.7906 | 0.0010 | fixed |
| Random effect mother ID | 0.0256 | 0.0959 | 0.1446 | 0.0022 | 0.4674 | 430.3779 | NA | random |
| Random effect year | 0.3879 | 0.4220 | 0.3521 | 0.0032 | 0.9543 | 624.0664 | NA | random |

**Table S3.** Full model results of the analysis of salmon abundance in current year (“Current salmon”) and in the previous year (“Previous salmon) with a minimum sighting threshold of five per year. For each effect we give the posterior distribution (PD) mode, median, and mean, the lower and upper 95% credible intervals (CI), and the effective sample size. For the fixed effects we also give the pMCMC values.

| Variable | PD mode | PD median | PD mean | Lower 95% CI | Upper 95% CI | Effective sample size | pMCMC | Effect type |
| --- | --- | --- | --- | --- | --- | --- | --- | --- |
| Intercept | -1.5300 | -1.4495 | -1.3679 | -1.8499 | -0.9623 | 706.8619 | 0.0010 | fixed |
| Current salmon | -0.2088 | -0.1532 | -0.1312 | -0.6362 | 0.3326 | 1000.0000 | 0.5680 | fixed |
| Previous salmon | -0.0743 | -0.1331 | -0.1918 | -0.7213 | 0.2725 | 769.4339 | 0.4160 | fixed |
| If the female had a calf in the last two years | -2.0460 | -2.1312 | -2.2679 | -3.0907 | -1.4646 | 653.5009 | 0.0010 | fixed |
| Random effect mother ID | 0.0256 | 0.0959 | 0.2937 | 0.0025 | 0.9809 | 421.2695 | NA | random |
| Random effect year | 0.3879 | 0.4220 | 0.4747 | 0.0045 | 1.2353 | 504.7265 | NA | random |

**Table S4.** Full model results of the analysis of the fitness benefits of mean and plasticity in individual strength at the monthly scale. This was a bivariate model with response variables of strength and calving success. Strength had random slopes and random intercepts and so we estimated the among-individual covariances between these two terms and between each and individual calving success. For each effect we give the posterior distribution (PD) mode, median, and mean; the lower and upper 95% credible intervals (CI); and the effective sample size. For the fixed effects we also give the pMCMC values.

| Variable | PD mode | PD median | PD mean | Lower 95% CI | Upper 95% CI | Effective sample size | pMCMC | Effect type |
| --- | --- | --- | --- | --- | --- | --- | --- | --- |
| Intercept – Strength | 7.0662 | 6.8380 | 6.8522 | 5.2229 | 8.3900 | 1000.0000 | 0.0010 | fixed |
| Intercept – Calving success | 0.1756 | 0.1732 | 0.1729 | 0.1475 | 0.1964 | 1000.0000 | 0.0010 | fixed |
| Salmon abundance on strength | 0.1349 | 0.2638 | 0.2616 | -0.2260 | 0.6692 | 1000.0000 | 0.2360 | fixed |
| Random effect dolphin ID - Strength | 2.2940 | 2.6148 | 2.7848 | 1.2007 | 4.7068 | 1000.0000 | NA | random |
| Random effect dolphin ID – Strength slope | 0.0036 | 0.1183 | 0.1762 | 0.0000 | 0.5261 | 1000.0000 | NA | random |
| Random effect dolphin ID – Calving success | 0.0090 | 0.0098 | 0.0100 | 0.0064 | 0.0136 | 294.5699 | NA | random |
| Covariance – Strength intercept & Calving success | 0.0233 | 0.0279 | 0.0282 | -0.0273 | 0.0855 | 1000.0000 | NA | random |
| Covariance – Strength intercept & slope | 0.0003 | -0.1202 | -0.1867 | -0.8776 | 0.2679 | 1000.0000 | NA | random |
| Covariance – Strength slope & Calving success | 0.0000 | -0.0054 | -0.0081 | -0.0345 | 0.0128 | 1000.0000 | NA | random |
| Random effect of year - Strength | 2.7622 | 2.8726 | 3.0686 | 1.3560 | 5.4746 | 1000.0000 | NA | random |
| Random effect of month - Strength | 0.7650 | 1.5046 | 2.5681 | 0.1189 | 7.1535 | 1000.0000 | NA | random |
| Residual variance - Strength | 8.8456 | 9.1507 | 9.1780 | 8.1877 | 10.2215 | 1440.4670 | NA | residual |

**Table S5.** Full model results of the analysis of the fitness benefits of mean and plasticity in individual closeness at the monthly scale. This was a bivariate model with response variables of closeness and calving success. Closeness had random slopes and random intercepts and so we estimated the covariances between these two terms and between each and individual calving success. For each effect we give the posterior distribution (PD) mode, median, and mean, the lower and upper 95% credible intervals (CI), and the effective sample size. For the fixed effects we also give the pMCMC values.

| Variable | PD mode | PD median | PD mean | Lower 95% CI | Upper 95% CI | Effective sample size | pMCMC | Effect type |
| --- | --- | --- | --- | --- | --- | --- | --- | --- |
| Intercept – Closeness | 0.5663 | 0.5467 | 0.5460 | 0.4464 | 0.6479 | 1000.0000 | 0.0010 | fixed |
| Intercept – Calving success | 0.1670 | 0.1722 | 0.1722 | 0.1473 | 0.1971 | 1000.0000 | 0.0010 | fixed |
| Salmon abundance on Closeness | 0.0010 | -0.0022 | -0.0025 | -0.0274 | 0.0187 | 678.4553 | 0.8720 | fixed |
| Random effect dolphin ID - Closeness | 0.0010 | 0.0016 | 0.0017 | 0.0000 | 0.0037 | 1000.0000 | 0.0010 | random |
| Random effect dolphin ID – Closeness slope | 0.0000 | 0.0002 | 0.0003 | 0.0000 | 0.0011 | 1000.0000 | NA | random |
| Random effect dolphin ID – Calving success | 0.0092 | 0.0094 | 0.0096 | 0.0067 | 0.0135 | 619.9407 | NA | random |
| Covariance – Closeness intercept & Calving success | 0.0009 | 0.0009 | 0.0010 | -0.0008 | 0.0030 | 1000.0000 | NA | random |
| Covariance – Closeness intercept & slope | -0.0000 | -0.0002 | -0.0003 | -0.0010 | 0.0002 | 1000.0000 | NA | random |
| Covariance –Closeness slope & Calving success | 0.0000 | -0.0003 | -0.0004 | -0.0018 | 0.0006 | 814.4118 | NA | random |
| Random effect of year - Closeness | 0.0113 | 0.0118 | 0.0126 | 0.0054 | 0.0210 | 1000.0000 | NA | random |
| Random effect of month - Closeness | 0.0040 | 0.0062 | 0.0125 | 0.0013 | 0.0324 | 819.9268 | NA | random |
| Residual variance - Closeness | 0.0305 | 0.0302 | 0.0302 | 0.0270 | 0.0335 | 1000.0000 | NA | residual |

**Table S6.** Full model results of the analysis of the fitness benefits of mean and plasticity in individual strength at the yearly scale. This was a bivariate model with response variables of strength and calving success. Strength had random slopes and random intercepts and so we estimated the covariances between these two terms and between each and individual calving success. For each effect we give the posterior distribution (PD) mode, median, and mean, the lower and upper 95% credible intervals (CI), and the effective sample size. For the fixed effects we also give the pMCMC values.

| Variable | PD mode | PD median | PD mean | Lower 95% CI | Upper 95% CI | Effective sample size | pMCMC | Effect type |
| --- | --- | --- | --- | --- | --- | --- | --- | --- |
| Intercept – Strength | 5.3990 | 5.4112 | 5.4102 | 4.7520 | 6.0690 | 1000.0000 | 0.0010 | fixed |
| Intercept – Calving success | 0.1655 | 0.1688 | 0.1691 | 0.1442 | 0.1903 | 864.9448 | 0.0010 | fixed |
| Salmon abundance on strength | 0.2999 | 0.2524 | 0.2511 | -0.2675 | 0.8024 | 826.3697 | 0.3360 | fixed |
| Random effect dolphin ID - Strength | 1.7355 | 1.7979 | 1.8505 | 1.0588 | 2.8285 | 1133.7821 | NA | random |
| Random effect dolphin ID – Strength slope | 0.0002 | 0.0160 | 0.0320 | 0.0000 | 0.1170 | 1000.0000 | NA | random |
| Random effect dolphin ID – Calving success | 0.0082 | 0.0093 | 0.0095 | 0.0065 | 0.0132 | 760.5222 | NA | random |
| Covariance – Strength intercept & Calving success | 0.0326 | 0.0314 | 0.0321 | -0.0059 | 0.0762 | 903.9309 | NA | random |
| Covariance – Strength intercept & slope | 0.0000 | -0.0200 | -0.0393 | -0.2474 | 0.1063 | 1000.0000 | NA | random |
| Covariance – Strength slope & Calving success | -0.0001 | -0.0002 | -0.0010 | -0.0161 | 0.0121 | 1000.0000 | NA | random |
| Random effect of year - Strength | 2.4168 | 2.4413 | 2.5356 | 1.3618 | 4.1571 | 1000.0000 | NA | random |
| Residual variance - Strength | 4.1281 | 4.1775 | 4.1915 | 3.6751 | 4.6948 | 1273.3871 | NA | residual |

**Table S7.** Full model results of the analysis of the fitness benefits of mean and plasticity in individual closeness at the yearly scale. This was a bivariate model with response variables of closeness and calving success. Closeness had random slopes and random intercepts and so we estimated the covariances between these two terms and between each and individual calving success. For each effect we give the posterior distribution (PD) mode, median, and mean, the lower and upper 95% credible intervals (CI), and the effective sample size. For the fixed effects we also give the pMCMC values.

| Variable | PD mode | PD median | PD mean | Lower 95% CI | Upper 95% CI | Effective sample size | pMCMC | Effect type |
| --- | --- | --- | --- | --- | --- | --- | --- | --- |
| Intercept – Closeness | 0.6286 | 0.6218 | 0.6197 | 0.5412 | 0.6916 | 1000.0000 | 0.0010 | fixed |
| Intercept – Calving success | 0.1738 | 0.1691 | 0.1687 | 0.1463 | 0.1927 | 1000.0000 | 0.0010 | fixed |
| Salmon abundance on Closeness | -0.0172 | -0.0164 | -0.0160 | -0.0824 | 0.0487 | 1000.0000 | 0.6320 | fixed |
| Random effect dolphin ID - Closeness | 0.0091 | 0.0096 | 0.0098 | 0.0051 | 0.0141 | 1000.0000 | NA | random |
| Random effect dolphin ID – Closeness slope | 0.0001 | 0.0003 | 0.0004 | 0.0000 | 0.0009 | 1000.0000 | NA | random |
| Random effect dolphin ID – Calving success | 0.0087 | 0.0093 | 0.0094 | 0.0062 | 0.0124 | 505.9510 | NA | random |
| Covariance – Closeness intercept & Calving success | 0.0026 | 0.0034 | 0.0035 | 0.0005 | 0.0068 | 1000.0000 | NA | random |
| Covariance – Closeness intercept & slope | 0.0020 | 0.0022 | 0.0023 | -0.0004 | 0.0050 | 1000.0000 | NA | random |
| Covariance –Closeness slope & Calving success | -0.0003 | -0.0005 | -0.0006 | -0.0019 | 0.0006 | 1000.0000 | NA | random |
| Random effect of year - Closeness | 0.0347 | 0.0411 | 0.0428 | 0.0232 | 0.0643 | 772.9455 | NA | random |
| Residual variance - Closeness | 0.0136 | 0.0139 | 0.0139 | 0.0123 | 0.0155 | 1000.0000 | NA | residual |
